# Nanoscale numerical simulations explain apparently opposing experimental findings on ephaptic coupling

**DOI:** 10.64898/2026.08.11.744093

**Authors:** Karoline Horgmo Jæger, Aslak Tveito

**Affiliations:** Simula Research Laboratory, Norway

## Abstract

A classical study found no excitation transfer when isolated cardiomyocytes were placed side by side, whereas a recent paper reported action potential transfer in carrdiomyocytes placed end to end. We use nanoscale numerical simulations based on the full Poisson–Nernst–Planck equations to investigate whether these apparently opposing observations can be explained by the different geometrical configurations. The computations show that in the end-to-end configuration, ephaptic coupling occurs when the intercellular cleft is sufficiently narrow and a sufficiently large fraction of the sodium channels is localized at the intercalated disc. Coupling is strengthened when the sodium channels are concentrated in fewer clusters and when ionic diffusion within the cleft is reduced. Under these conditions, excitation transfer occurs on a timescale consistent with rapid cell-to-cell activation. Conduction depends biphasically on cleft width and terminates abruptly beyond a critical width. Localization of potassium channels at the intercalated disc has only a moderate effect, whereas gap junctions substantially improve conduction and reduce the relative contribution of ephaptic coupling. In the side-by-side configuration, excitation transfer does not occur under physiological conditions and requires highly flattened cells, minimal separation, and unrealistically strong sodium-channel clustering. The different outcomes of the side-by-side and end-to-end experiments can therefore be explained by the fundamentally different geometrical conditions for ephaptic coupling.

## 1 Introduction

A classical study [1] found no excitation transfer when isolated cardiomyocytes were placed side by side, whereas a recent paper [2] reported action potential (AP) transfer in cells placed end to end. These apparently opposing findings sharpen a longstanding question: can one cell excite its neighbor through the extracellular space alone, without direct coupling through gap junctions or chemical synapses? This question has been examined extensively in both cardiology, [3, 4, 5], and neuroscience, [6, 7, 8, 9], using both experimental, [1, 10, 11, 2], and modeling approaches, [12, 13, 14, 15, 16, 17, 18].

An early and influential answer was provided by Weingart and Maurer in 1988, [1]. They isolated pairs of cardiomyocytes, placed them side by side, and summarized their observations as follows: *Two single myocytes, gently pushed together, neither showed electrotonic interaction nor impulse transfer, thus rendering unlikely the possibility of an ephaptic signal transmission*.

Despite this conclusion, the question remained active, and investigations continued in both cardiology and neuroscience. In 2024, Waaben et al., [2], revisited the experiment using isolated cardiomyocyte pairs. In contrast to the side-by-side configuration considered by Weingart and Maurer, the cells were placed end to end, such that the sodium-channel-rich membranes of the intercalated discs (IDs) faced each other across a narrow extracellular cleft. They concluded: *In summary, this manuscript demonstrates that cardiomyocytes can activate each other by ephaptic transmission alone, substantiates that ephaptic transmission contributes to AP transmission in native cardiomyocyte pairs and for the first time demonstrates that even isolated cardiomyocyte pairs retain shielded domains, a prerequisite for ephaptic transmission*.

The two studies therefore reached apparently opposing conclusions under fundamentally different geometrical conditions. In the side-by-side configuration, the closely apposed membranes are located along the lateral surfaces of the cells, whereas in the end-to-end configuration, the sodium-channel-rich membranes of the IDs face each other across a narrow cleft. This difference suggests that cell geometry, membrane separation, and the spatial organization of ion channels may determine whether ephaptic coupling (EpC) is possible.

Additional experimental studies have identified structural and functional conditions that may support EpC in cardiac tissue. At the ID, sodium channels (NaChs) are concentrated within specialized nanodomains adjacent to gap junctions (GJs), where opposing membranes may be separated by distances on the nanometer scale. This perinexal region has therefore been proposed as a structural substrate for ephaptic transmission, while also providing a close spatial association between ephaptic and gap-junctional coupling, [4]. Furthermore, using osmotic agents to alter perinexal width without measurable changes in peak sodium current, Cx43 expression, or GJ conductance, it has been shown that cardiac conduction velocity (CV) depends biphasically on perinexal separation, indicating that the width of the extracellular cleft influences conduction even when GJs are present, [11]. Recent experiments on isolated cardiomyocyte pairs have sought to separate the contributions of GJ coupling and EpC by varying GJ conductance, extracellular sodium concentration, and perinexal width. These results indicate that EpC can provide substantial support for intercellular activation, particularly when GJ coupling is reduced, while the two mechanisms may operate cooperatively under physiological conditions. Together, these findings support a mixed mechanism of cardiac conduction whose relative contributions depend on the structure and ionic environment of the ID, [5].

Computational studies have played a central role in establishing the conditions under which EpC may contribute to cardiac conduction. Early circuit models showed that a negative extracellular potential in the intercellular cleft can activate the post-junctional membrane and that EpC and GJ coupling may act together, [19]. Subsequent cable-based models demonstrated that localization of NaChs at the ID can either slow conduction through self-attenuation or promote excitation transfer when GJ coupling is strongly reduced, [12]. Related models examined the influence of extracellular microdomains on transverse propagation, [20]. Spatially resolved finite element models later showed that clustering of NaChs strengthens ephaptic interactions, [21]. A cell-based model with explicitly represented intracellular and extracellular spaces further showed that sufficiently narrow clefts may support ephaptic excitation transfer in the absence of GJs, [22]. Models of cardiomyocyte pairs have also investigated how EpC and GJ coupling affect automaticity and excitation transfer, [17]. More recent electrodiffusion models, under the assumption of electroneutrality, have incorporated dynamic ionic concentrations and realistic ID nanostructures, identifying Na^+^ depletion, Na^+^ transfer, and cleft heterogeneity as additional modulators of EpC, [23]. Finally, recent cable simulations have examined how EpC modifies the source–sink relation and its dependence on cleft width, channel localization, and GJ conductance, [24].

A previous study applied the full Poisson–Nernst–Planck (PNP) equations to a small part of the ID between two cardiomyocytes to examine nanoscale effects while allowing for non-electroneutral conditions in the narrow space between the cell membranes, [18]. The severe computational requirements restricted the simulations to a small spatial domain and a short time interval, but the results indicated that the extracellular potential generated by NaCh activation could be sufficiently large to support EpC. The numerical method has since been substantially improved by solving the concentration equations and Poisson equation in a coupled implicit scheme, [25, 26]. This removes the severe time-step restriction associated with the earlier operator-splitting method, as subsequently analyzed in [27]. These developments make it possible to accurately simulate the adjoining halves of two cardiomyocytes throughout excitation transfer.

We use this approach to investigate whether the apparently opposing outcomes of the side-by-side and end-to-end experiments can be explained by their different geometrical configurations. We first determine the conditions under which EpC can support excitation transfer between end-to-end cardiomyocytes. We find that EpC requires a sufficiently narrow cleft and a sufficiently large fraction of NaChs at the ID, and is strengthened when the channels are concentrated in fewer clusters or when cleft diffusion is reduced. The activation delay depends biphasically on cleft width and increases abruptly as the system approaches conduction block. Potassium channels at the ID have a moderate effect, whereas GJs substantially improve conduction and reduce the importance of EpC under normal coupling conditions.

We then investigate the side-by-side configuration. EpC does not occur for standard round cell geometries, even under restricted diffusion. When flattened cell morphologies are combined with nanometer-scale separations, excitation transfer only emerges with unrealistically strong localized NaCh clustering. The simulations therefore explain why EpC may occur in the end-to-end configuration while remaining absent in the side-by-side arrangement, thereby reconciling the classical observations of Weingart and Maurer, [1], with the recent findings of Waaben et al., [2].

## 2 Results

To investigate the mechanisms and requirements for ephaptic coupling between cardiomyocytes, we perform a series of simulations using the PNP model. Unless otherwise stated, the simulations consider pure EpC, with no gap junctions connecting the cells. The majority of the simulations examine two cardiomyocytes arranged end-to-end like illustrated in Figure 1, whereas the final set of simulations considers a side-by-side arrangement like illustrated in Figure 2. We explore the effects of several factors that are expected to influence EpC, including the cleft width, the concentration and clustering of NaChs at the ID, and the diffusion properties of the extracellular cleft.

**Figure 1:**
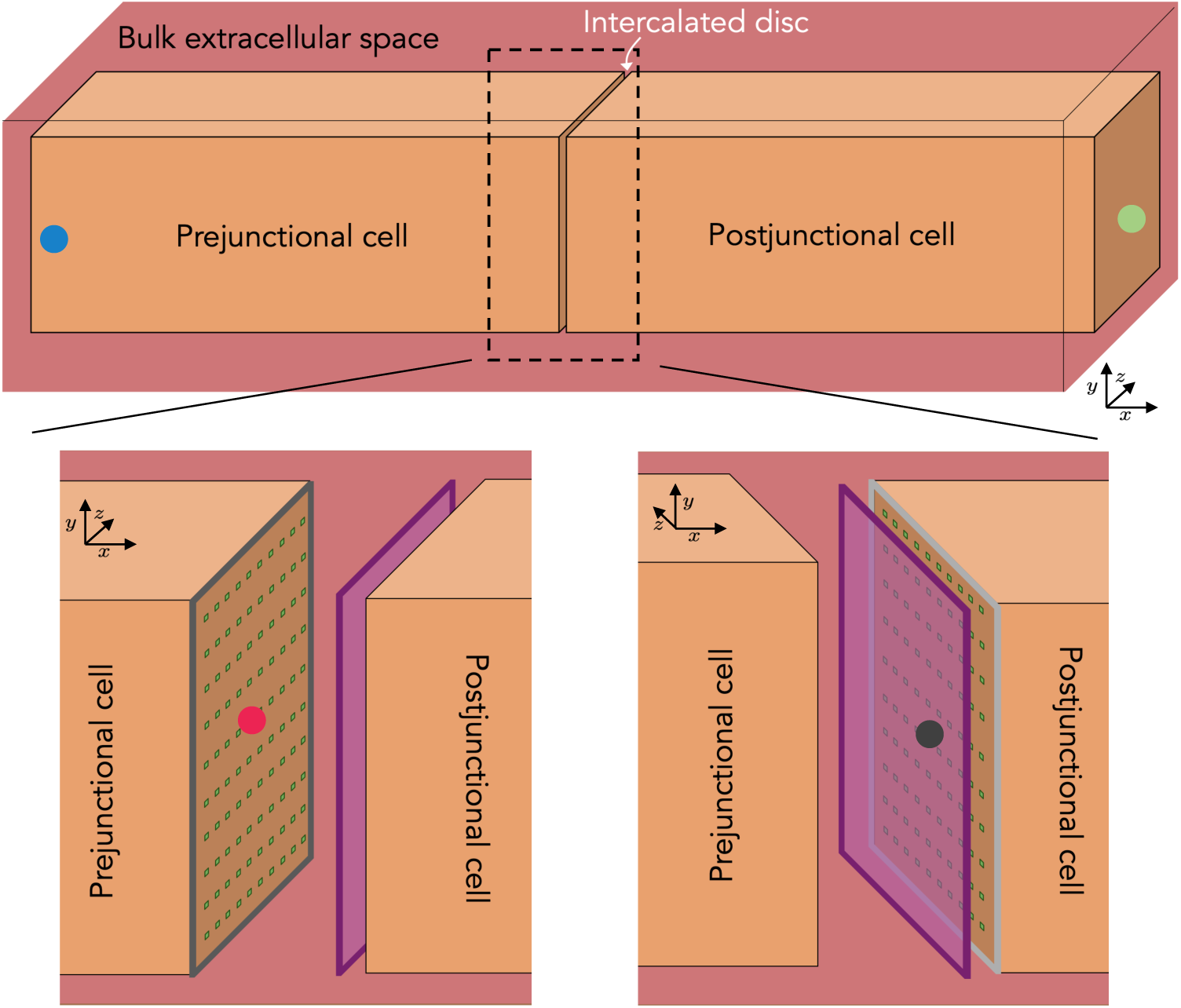
End-to-end arrangement of two cardiomyocytes. The upper panel shows the full computational domain with a blue and a green dot marking measurement points in the prejunctional and postjunctional cardiomyocytes, respectively. In the lower panel, we zoom in on the intercalated disc between the cells and mark measurement points on the membrane of the two cells and a measurement plane between the cells.

**Figure 2:**
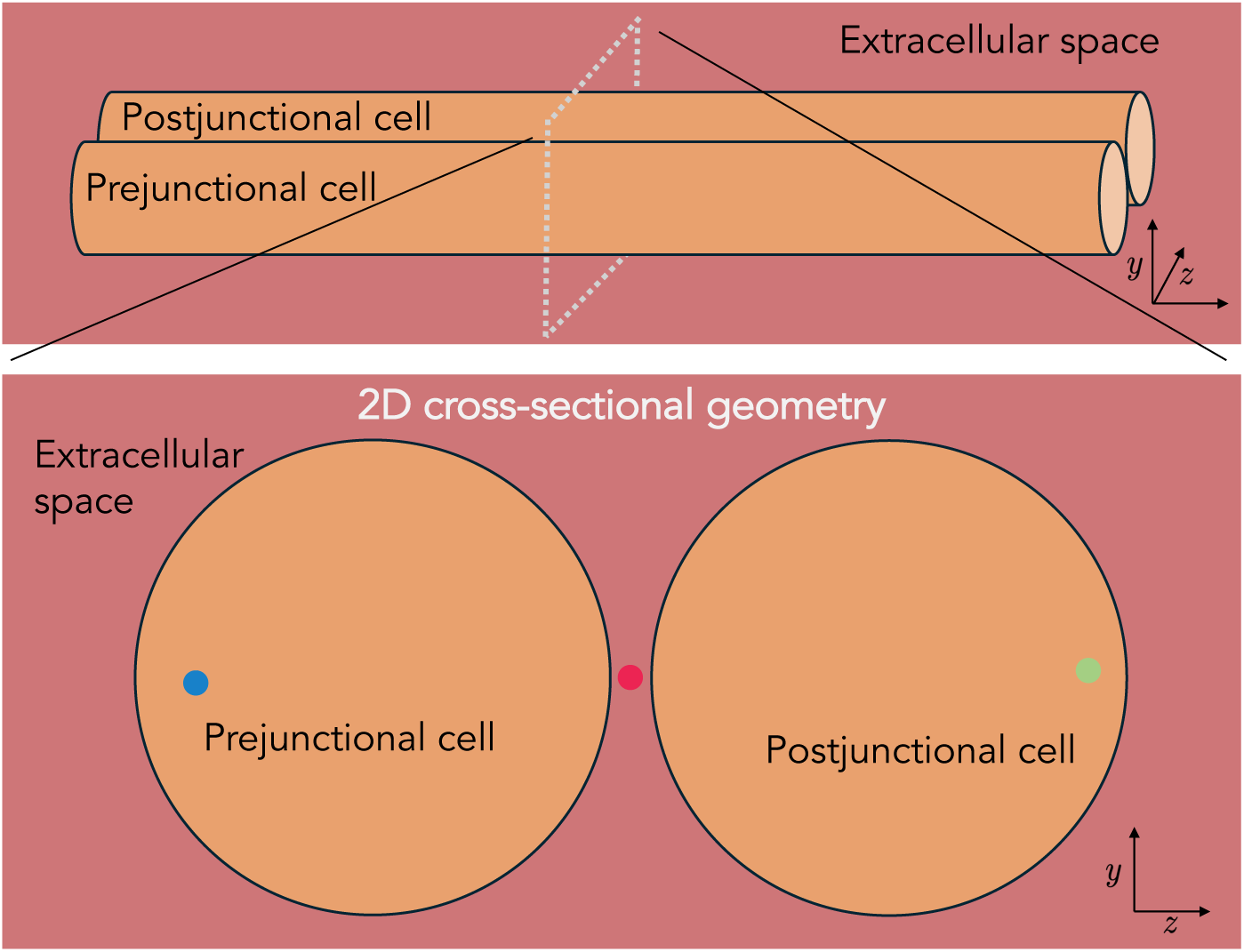
Side-by-side arrangement of two cardiomyocytes. The computational domain consists of a prejunctional and a postjunctional cell surrounded by an extracellular space. This computational domain is represented in 2D (*y, z*), assuming no spatial gradients in the *x*-direction, as illustrated in the lower panel. Measurement point for the prejunctional cell, postjunctional cell and the extracellular space between the cells are indicated by blue, green and red dots, respectively.

### 2.1 Concentration of sodium channels strengthens ephaptic coupling

We begin by investigating how the concentration and clustering of NaChs at the ID influence EpC. To this end, we vary the fraction of the cardiomyocyte’s total NaCh population located at the ID (*f*_Na_) from 50% to 100%. We also examine the effect of channel clustering by distributing the ID-localized NaChs into either 16 or 100 equally sized clusters, as described in the Methods section. For the case in which 80% of the NaChs are located at the ID, these configurations correspond to cluster widths of 0.6 *µ*m and 0.24 *µ*m for 16 and 100 clusters, respectively, which are comparable to experimentally observed NaCh cluster sizes [28, 29, 30].

Figure 3 shows the occurrence of EpC between the cardiomyocytes, in the absence of gap junctions, as a function of ID NaCh concentration and cleft width. We find that ephaptic coupling becomes more likely as a larger fraction of NaChs is concentrated at the ID, the extracellular cleft becomes narrower, or the NaCh clusters become larger and fewer.

**Figure 3:**
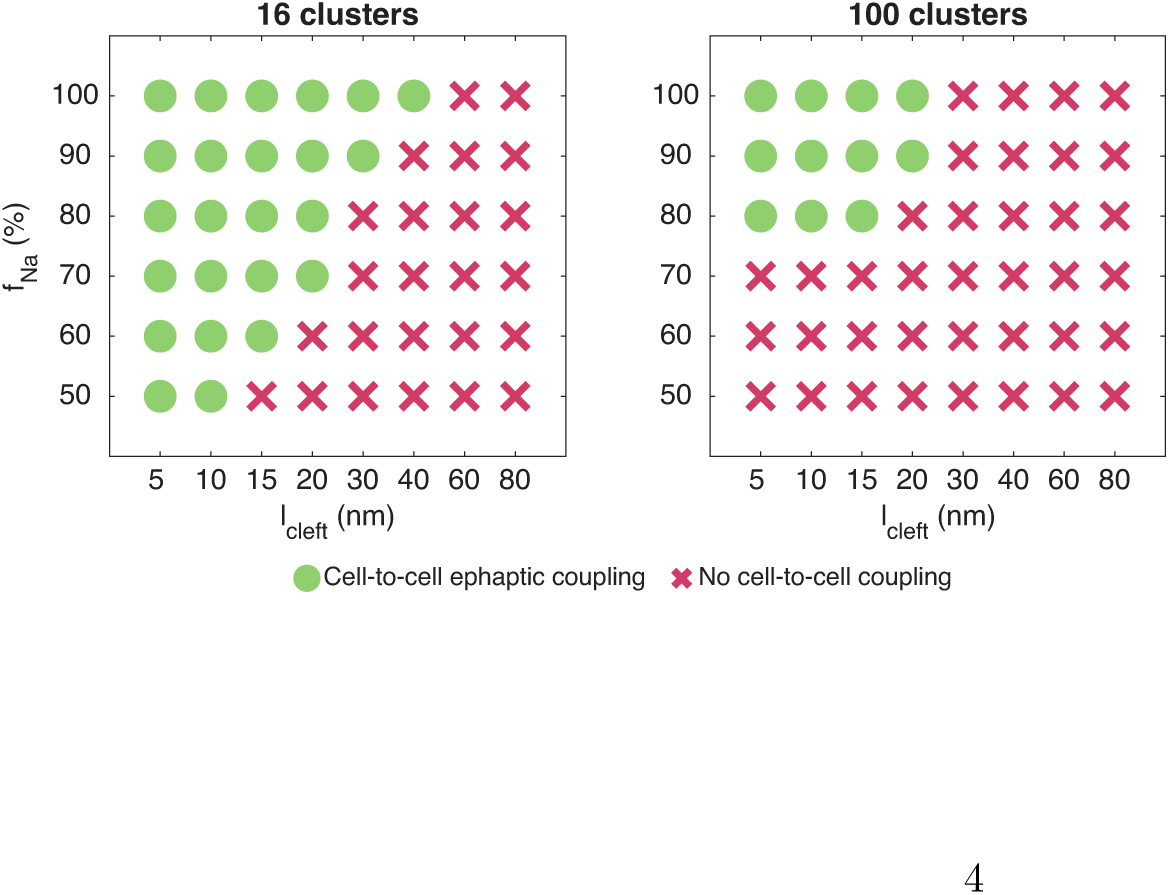
Occurrence of action potential transfer via ephaptic coupling for varying concentrations of sodium channels at the intercalated disc and cleft widths. We consider two cardiomyocytes arranged end-to-end, like illustrated in Figure 1. Here, *f*_Na_ denotes the fraction of the cell’s total sodium channels located at the intercalated disc, and *l*_cleft_ denotes the width of the extracellular cleft separating the cells. The diffusion coefficients in the extracellular cleft are divided by a factor *κ* = 6. Details of the simulation setup and the criterion used to determine successful coupling are provided in the Methods section.

In Figures 4–6, we show time-dependent solutions evaluated at the points and planes defined in Figure 1 for selected parameter combinations. Figure 4 considers the case *f*_Na_ = 80%, *l*_cleft_ = 15 nm, and 16 sodium channel clusters. In this configuration, we observe that as the intracellular potential *φ_i_* of the prejunctional cell increases and its NaChs open, the extracellular potential in the cleft becomes negative. This, in turn, increases the postjunctional membrane potential *v* = *φ_i_ − φ_e_*, leading to opening of NaChs in the postjunctional membrane, starting at the center of the cleft and subsequently spreading across the ID. The resulting NaCh activation causes an increased intracellular potential of the postjunctional cell. Figure 5 shows the corresponding dynamics in the case of 100 clusters. The overall sequence of events is similar to that in Figure 4, although activation of postjunctional NaChs occurs more slowly. Finally, Figure 6 shows the same configuration as in Figure 5, except that *f*_Na_ = 100% instead of *f*_Na_ = 80%. In this case, postjunctional NaCh activation occurs more rapidly than for *f*_Na_ = 80%.

**Figure 4:**
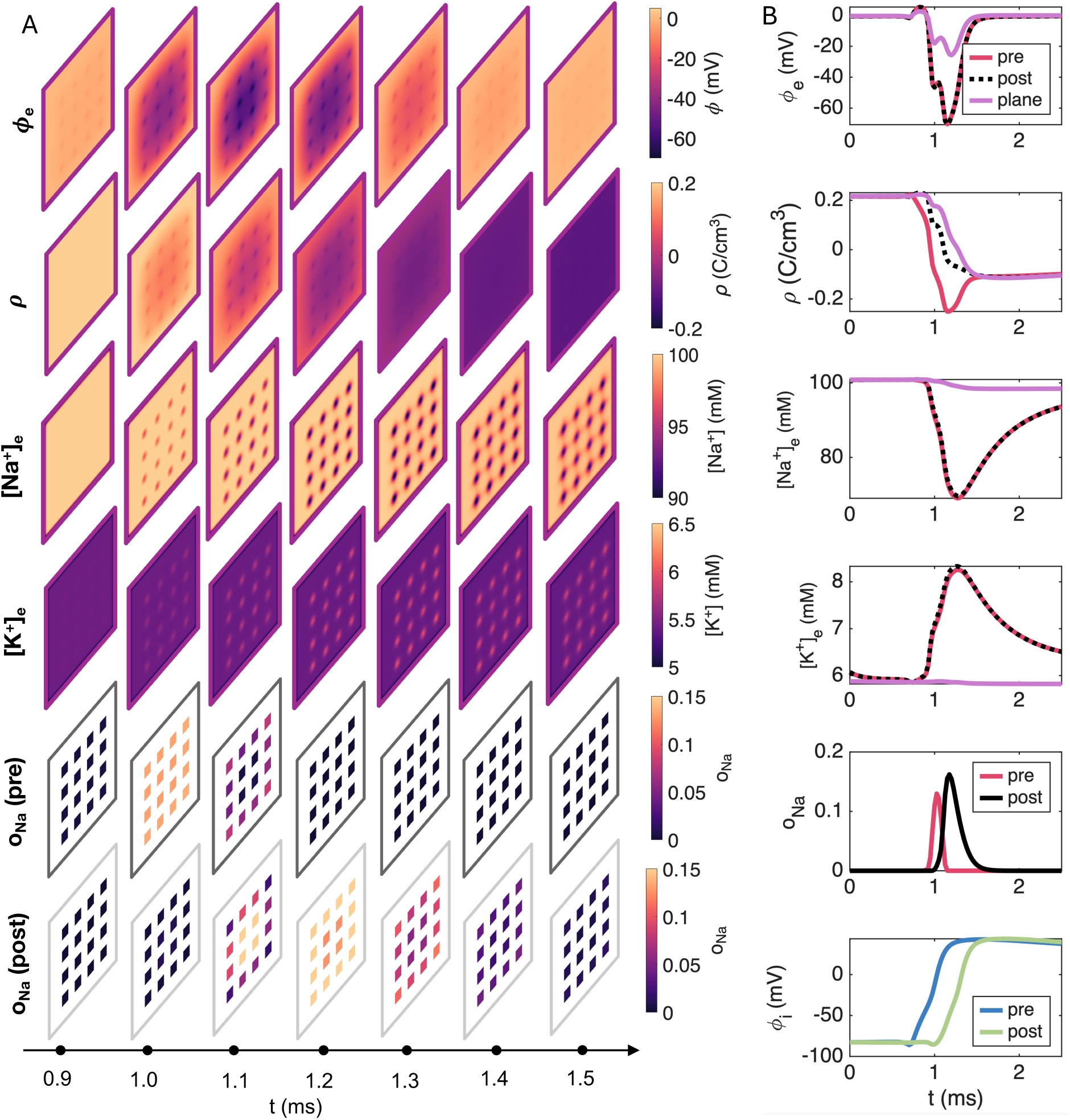
Simulated ephaptic coupling dynamics for a 15 nm wide cleft, 16 sodium channel clusters, *κ* = 6, and *f*_Na_= 80%. (A) Snapshots of the extracellular potential, *φ*, the net charge density, *ρ*, sodium concentration, [Na^+^], and potassium concentration, [K^+^] in a plane located 0.25 nm outside the postjunctional membrane (see Figure 1), as well as the sodium channel open probability, *o*_Na_, at the prejunctional and postjunctional ID membranes. (B) Time courses of *φ*, *ρ*, [Na^+^], [K^+^] and *o*_Na_ at the points defined in Figure 1. Spatial averages over the purple plane shown in Figure 1 are also included. Note that the red and black points are located at the centers of sodium channel clusters.

**Figure 5:**
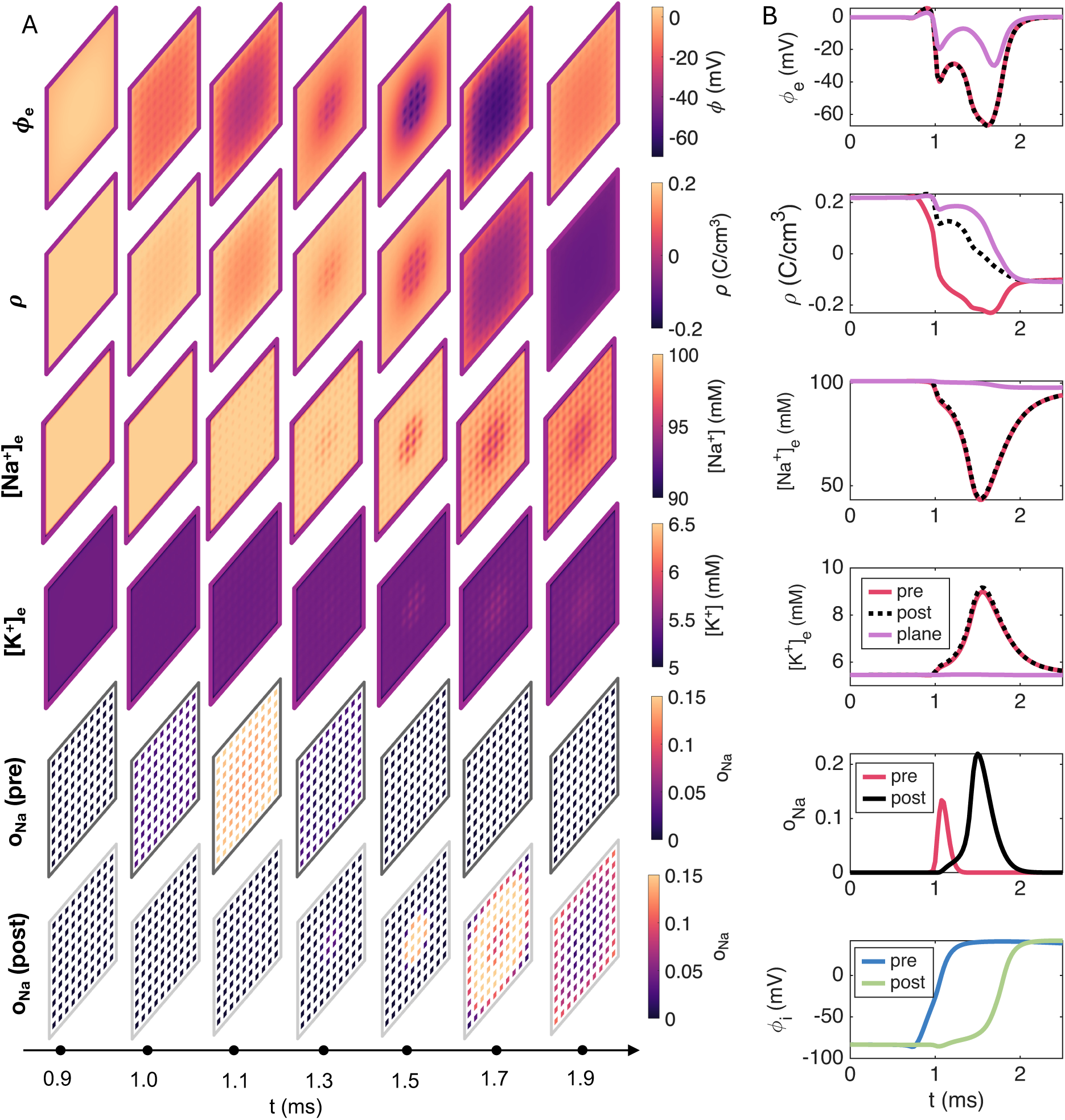
Simulated ephaptic coupling dynamics for a 15 nm wide cleft, *κ* = 6, 100 sodium channel clusters, and *f*_Na_= 80%. (A) Snapshots of the extracellular potential, *φ*, the net charge density, *ρ*, sodium concentration, [Na^+^], and potassium concentration, [K^+^] in a plane located 0.25 nm outside the postjunctional membrane (see Figure 1), as well as the sodium channel open probability, *o*_Na_, at the prejunctional and postjunctional ID membranes. (B) Time courses of *φ*, *ρ*, [Na^+^], [K^+^] and *o*_Na_ at the points defined in Figure 1. Spatial averages over the purple plane shown in Figure 1 are also included. Note that the red and black points are located at the centers of sodium channel clusters.

**Figure 6:**
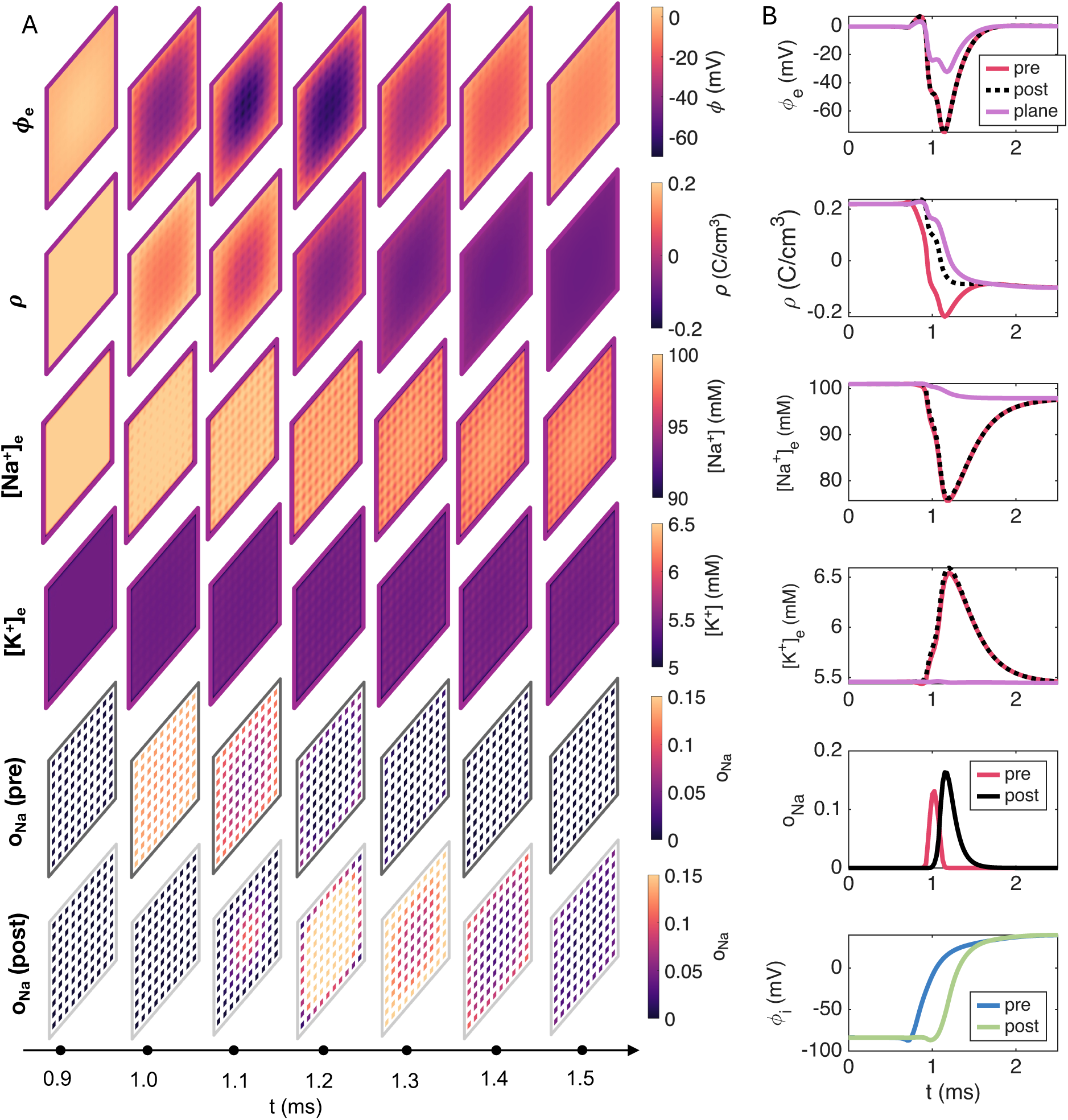
Simulated ephaptic coupling dynamics for a 15 nm wide cleft, *κ* = 6, 100 sodium channel clusters, and *f*_Na_= 100%. (A) Snapshots of the extracellular potential, *φ*, the net charge density, *ρ*, sodium concentration, [Na^+^], and potassium concentration, [K^+^] in a plane located 0.25 nm outside the postjunctional membrane (see Figure 1), as well as the sodium channel open probability, *o*_Na_, at the prejunctional and postjunctional ID membranes. (B) Time courses of *φ*, *ρ*, [Na^+^], [K^+^] and *o*_Na_ at the points defined in Figure 1. Spatial averages over the purple plane shown in Figure 1 are also included. Note that the red and black points are located at the centers of sodium channel clusters.

### 2.2 Decreasing cleft diffusion strengthens ephaptic coupling

The narrow extracellular cleft may hinder ion transport relative to free solution. To assess the impact of such diffusion restrictions on EpC, we perform simulations with reduced ionic diffusion coefficients in the cleft. All simulations presented thus far use a diffusion reduction factor of *κ* = 6, such that the diffusion coefficients in the cleft are reduced by a factor of six relative to their bulk values. In Figure 7, we investigate how varying *κ* between 1 (no diffusion restriction) and 10 (strong diffusion restriction) affects the occurrence of action potential transfer through EpC. We find that EpC becomes more likely as *κ* increases, indicating that restricted ion diffusion within the cleft promotes ephaptic interactions between the cells.

**Figure 7:**
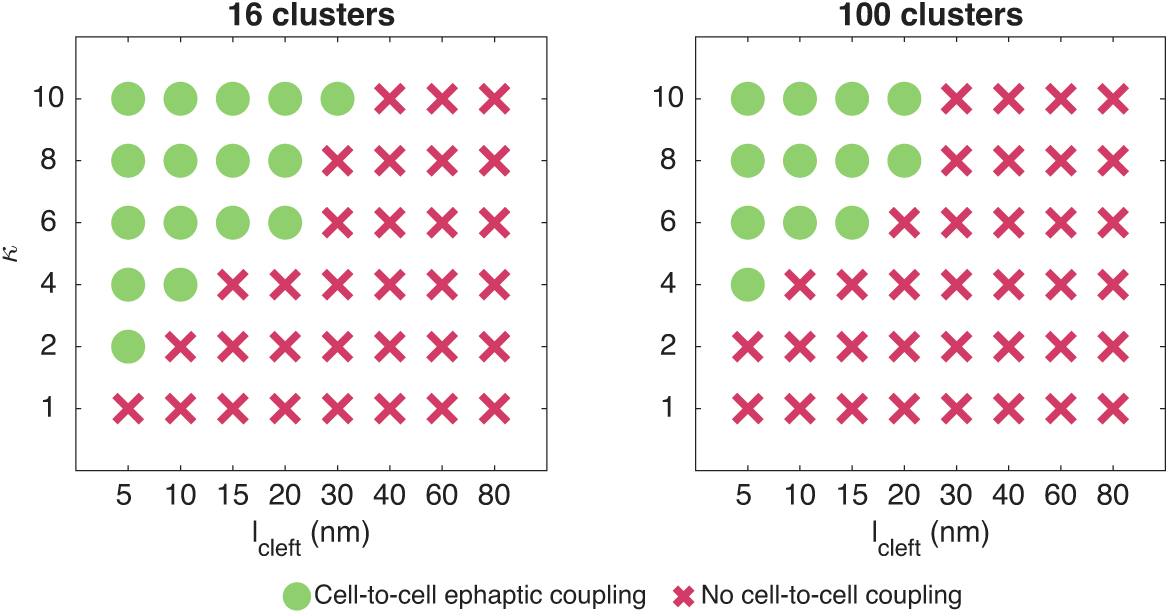
Occurrence of action potential transfer via ephaptic coupling for varying degrees of cleft diffusion reduction and varying cleft widths. We consider two cardiomyocytes arranged end-to-end, like illustrated in Figure 1. Here, *κ* denotes the factor by which the default diffusion coefficients in Table 1 are divided in the part of the domain representing extracellular space between the cells, and *l*_cleft_ denotes the cleft width. 80% of the sodium channels are located in the ID (*f*_Na_ = 80%). Details of the simulation setup

**Table 1:** PNP model parameter values used in the simulations. In the cell membrane, the diffusion coefficients are set to zero for all ions, and in the extracellular cleft, the default diffusion coefficients are divided by a factor *κ*.

| Parameter | Description | Value | Ref. |
| --- | --- | --- | --- |
| $F$ | Faraday's constant | 96485.3365 C/mol | [35] |
| $e$ | Elementary charge | $1.60217662 \cdot 10^{-19}$ C | [35] |
| $k_B$ | Boltzmann constant | $1.380649 \cdot 10^{-20}$ mJ/K | [35] |
| $T$ | Temperature | 310 K | |
| $\varepsilon_0$ | Vacuum permittivity | 8854 fF/m | [35] |
| $\varepsilon_m$ | Relative permittivity, $\varepsilon_r$, in the membrane | 2 | [36] |
| $\varepsilon_1$ | Relative permittivity, $\varepsilon_r$, elsewhere | 80 | [36] |
| $D_{\text{Na}^+}$ | Default diffusion coefficient for $\text{Na}^+$ | $1.33 \cdot 10^6$ nm<sup>2</sup>/ms | [37] |
| $D_{\text{K}^+}$ | Default diffusion coefficient for $\text{K}^+$ | $1.96 \cdot 10^6$ nm<sup>2</sup>/ms | [37] |
| $D_{\text{Ca}^{2+}}$ | Default diffusion coefficient for $\text{Ca}^{2+}$ | $0.71 \cdot 10^6$ nm<sup>2</sup>/ms | [37] |
| $D_{\text{Cl}^-}$ | Default diffusion coefficient for $\text{Cl}^-$ | $2.03 \cdot 10^6$ nm<sup>2</sup>/ms | [37] |
| $\kappa$ | Reduction factor for diffusion in the cleft | 6 | |
| $z_{\text{Na}^+}$ | Valence of $\text{Na}^+$ | 1 | |
| $z_{\text{K}^+}$ | Valence of $\text{K}^+$ | 1 | |
| $z_{\text{Ca}^{2+}}$ | Valence of $\text{Ca}^{2+}$ | 2 | |
| $z_{\text{Cl}^-}$ | Valence of $\text{Cl}^-$ | -1 | |

As a more detailed example, in Figure 8, we consider the same case as in Figure 4 except that diffusion is not reduced as much (*κ* = 2 instead of *κ* = 6). In this case, both the ionic concentration changes and the negative extracellular potential in the cleft are less pronounced. As a result, the extracellular potential generated by depolarization of the prejunctional cell does not become sufficiently negative to trigger NaCh opening in the postjunctional membrane.

**Figure 8:**
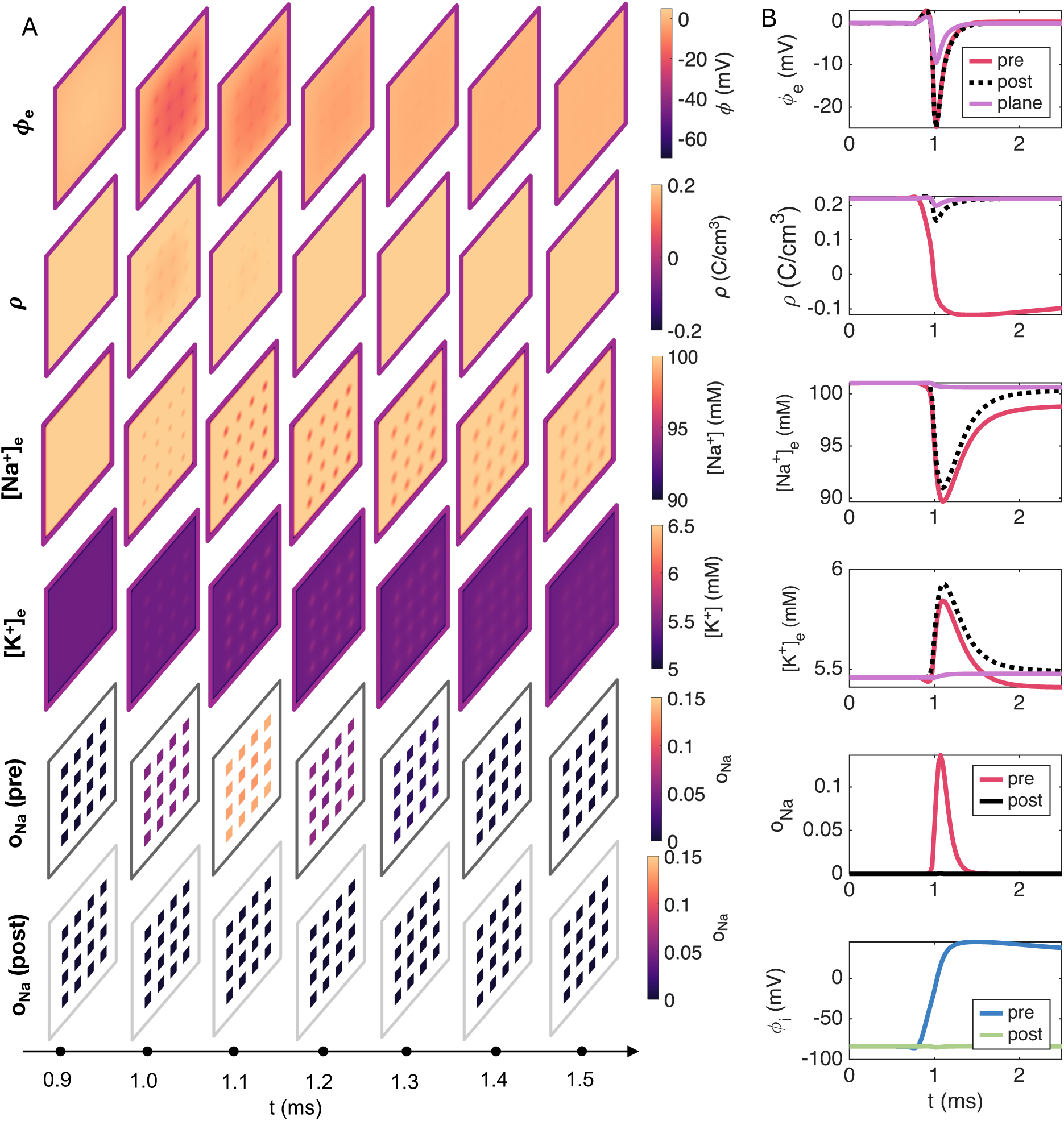
Simulated ephaptic coupling dynamics for a 15 nm wide cleft, 16 sodium channel clusters, *κ* = 2, and *f*_Na_= 80%. (A) Snapshots of the extracellular potential, *φ*, the net charge density, *ρ*, sodium concentration, [Na^+^], and potassium concentration, [K^+^] in a plane located 0.25 nm outside the postjunctional membrane (see Figure 1), as well as the sodium channel open probability, *o*_Na_, at the prejunctional and postjunctional ID membranes. (B) Time courses of *φ*, *ρ*, [Na^+^], [K^+^] and *o*_Na_ at the points defined in Figure 1. Spatial averages over the purple plane shown in Figure 1 are also included. Note that the red and black points are located at the centers of sodium channel clusters.

### 2.3 Conduction velocity varies with cleft width

So far, we have primarily focused on whether or not AP transfer through EpC occurs. In Figure 9, we further examine how the activation delay from the prejunctional to the postjunctional cell depends on the cleft width, the fraction of NaChs at the ID, and the reduction in cleft diffusion. Specifically, we report the activation delay, *τ*, and the corresponding CV for the cases in which AP transfer occurs in Figures 3 and 7. The observed activation delays range from 0.23 ms to 0.81 ms, corresponding to conduction velocities of approximately 12 cm/s to 44 cm/s, and we observe that there are abrupt transitions from conduction to conduction block. Moreover, for the case of 16 clusters, there is a biphasic dependence of the cleft width on the activation delays.

**Figure 9:**
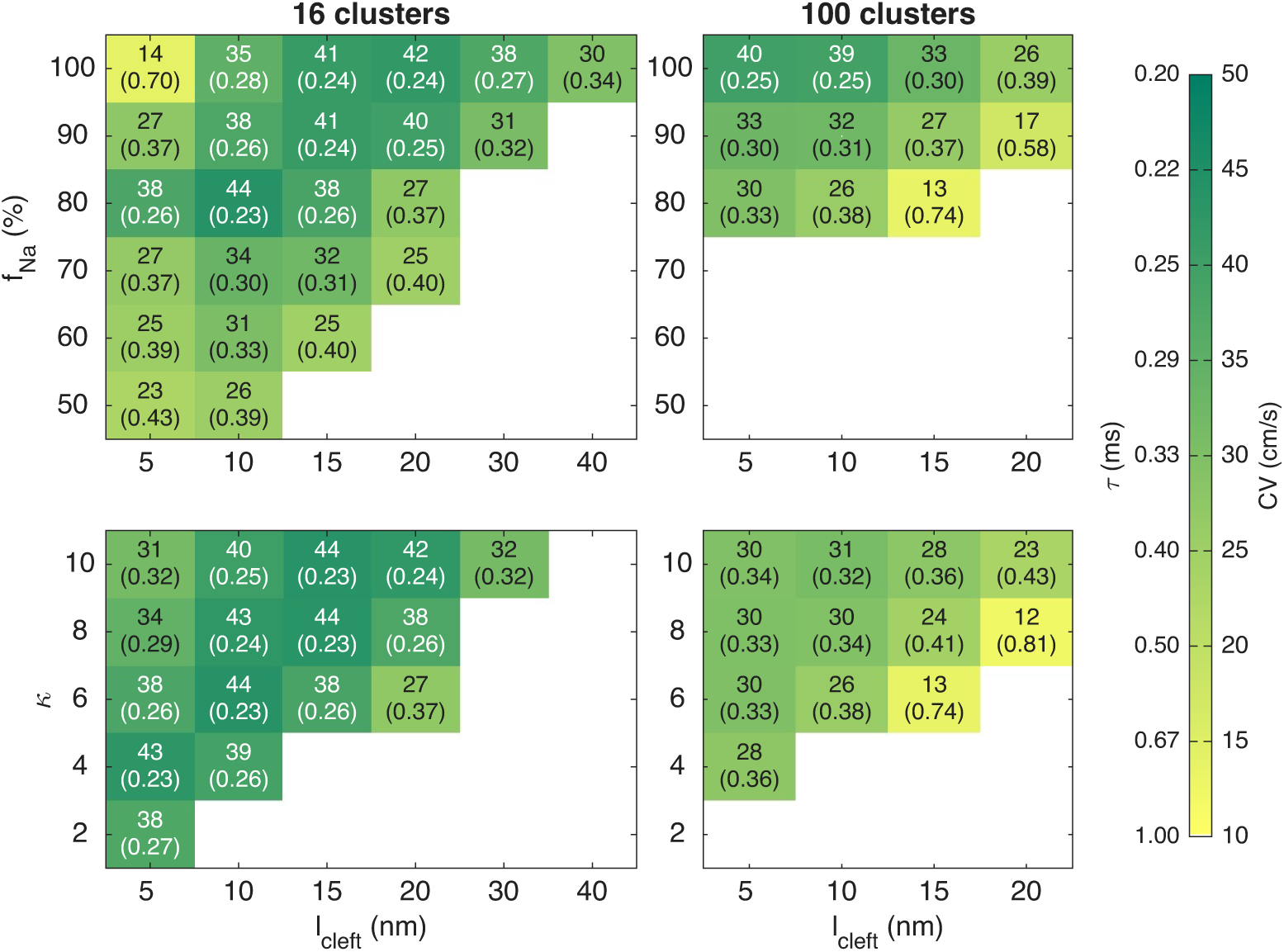
Conduction velocity (CV) and activation delay (*τ*) as functions of *l*_cleft_, *f*_Na_, and *κ*. The upper numbers report the CV and the lower numbers in parenthesis represents *τ*. We consider two cardiomyocytes arranged end-to-end, as illustrated in Figure 1. Here, *f*_Na_ denotes the fraction of the cell’s total sodium channels located at the intercalated disc, *κ* denotes the factor by which the diffusion coefficients are divided in the extracellular space between the cells, and *l*_cleft_ denotes the cleft width. In the upper panel *κ* = 6, while in the lower panel *f*_Na_ = 80%. Details of the simulation setup and the definition of *τ* are provided in the Methods section.

### 2.4 Potassium channels at the ID have a moderate effect on conduction

Experimental studies have shown that K_ir_2.1 potassium channels, which carry the *I*_K1_ current, are also enriched at the ID and within the perinexus [31, 32]. To investigate the potential impact of this localization on EpC, we perform simulations in which a fraction of the cell’s *I*_K1_ channels are co-localized with the NaCh clusters at the ID. Figure 10 shows the same simulation as in Figure 4, except that 40% of the *I*_K1_ channels are relocated to the IC. As expected, the potassium concentration in the vicinity of the channel clusters is considerably higher than in Figure 4. However, this redistribution has only a modest effect on AP transfer. The activation delay increases slightly from 0.26 ms to 0.29 ms, corresponding to a CV decrease from 38 cm/s to 34 cm/s.

**Figure 10:**
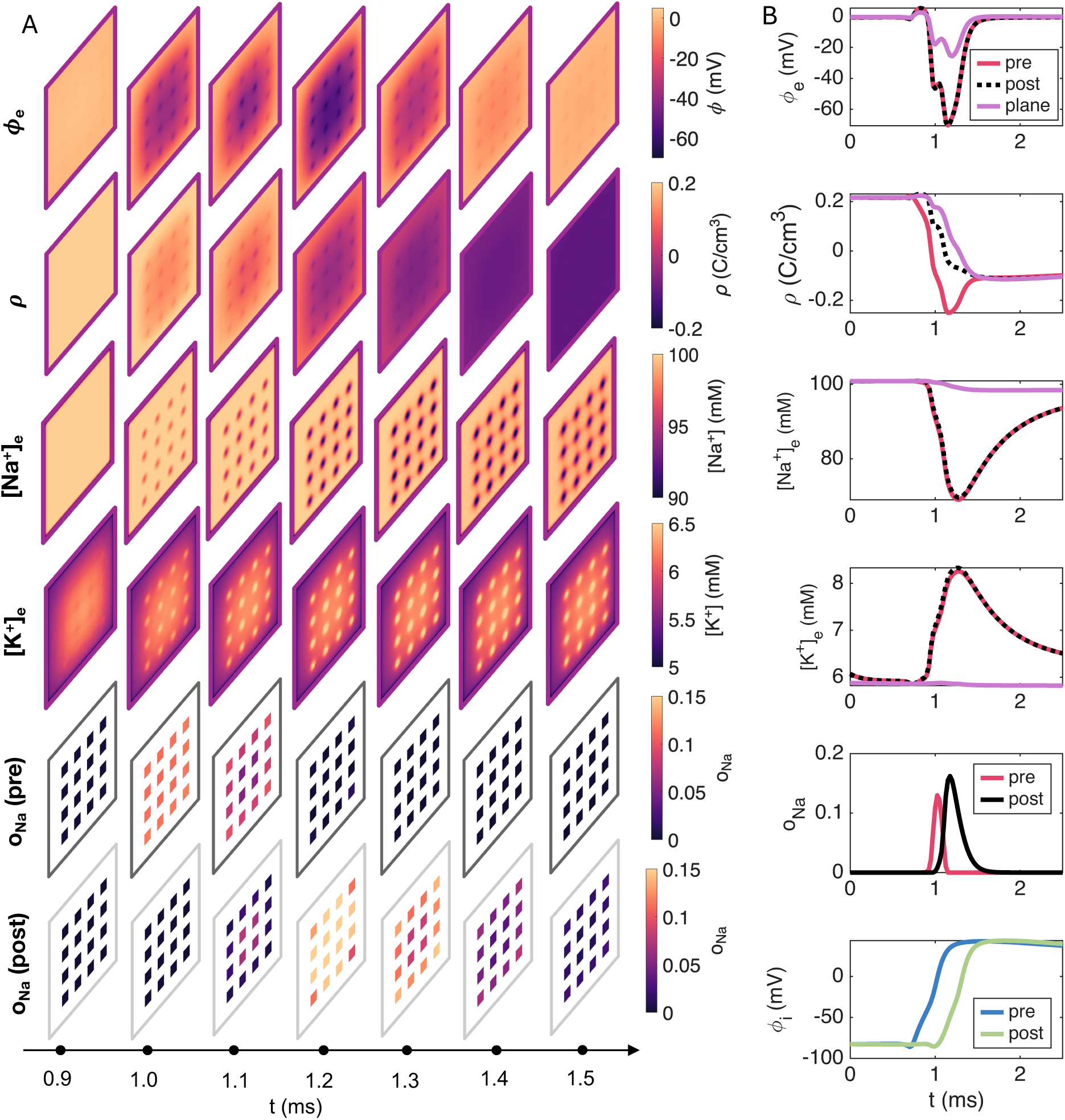
Simulated ephaptic coupling dynamics like in Figure 4, except that 40% of the K1 channels are located in the intercalated disc.

### 2.5 Gap junctions substantially improve conduction velocity

In the simulations presented thus far, we have considered pure EpC by omitting GJs between neighboring cells. In cardiac tissue, however, EpC and GJ-mediated coupling coexist, and ephaptic effects may therefore influence conduction even in the presence of GJs. To investigate this, Figure 11 shows the activation delay and CV for different cleft widths and GJ conductances. Throughout this analysis, we fix *f*_Na_ = 80% and *κ* = 6. As expected, increasing the GJ conductance decreases the activation delay and increases the conduction velocity compared with the case of pure EpC. Moreover, in contrast to the abrupt transition from conduction to block observed in the absence of GJs, the conduction velocity decreases more gradually as the GJ conductance is reduced. Moreover, the influence of cleft width persists over the entire range of GJ conductances considered, indicating that ephaptic interactions can modulate conduction even in the presence of GJs.

**Figure 11:**
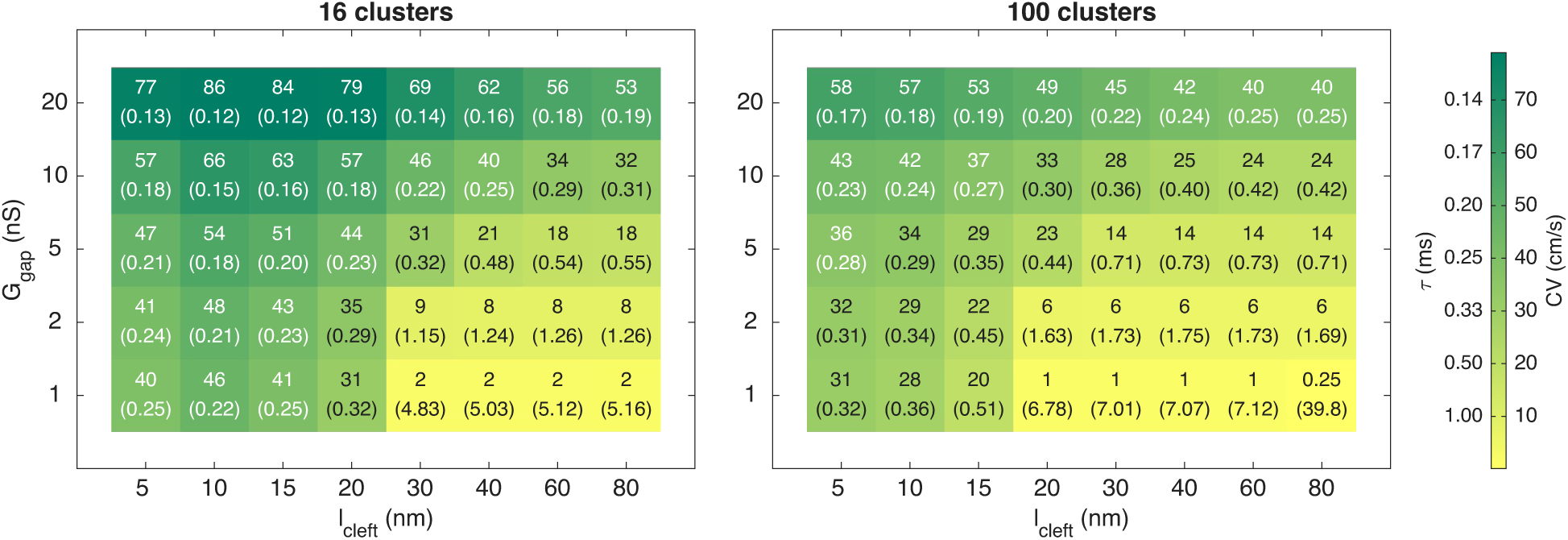
Conduction velocity (CV) and activation delay (*τ*) as functions of *l*_cleft_ and *G*_gap_ for simulations including gap junctions. The upper values report the CV (in cm/s) and the lower numbers in parenthesis represents *τ* (in ms). We consider two cardiomyocytes arranged end-to-end, as illustrated in Figure 1. Here, *G*gap denotes the total gap junction conductance and *l*_cleft_ denotes the cleft width. We use *f*_Na_ = 80% and *κ* = 6 Details of the simulation setup and the definition of *τ* are provided in the Methods section.

### 2.6 Side-by-side vs. end-to-end

In a final set of simulations, we investigate the potential for EpC in the side-by-side configuration. In the upper left panel of Figure 12, we report whether EpC occurs for intercellular distances in the range 2–80 nm for a default round cell cross-section. We also vary *f*_Na_, which in this case denotes the fraction of NaChs localized to a 0.5 *µ*m-wide membrane region adjacent to the region of closest separation between the cells. For a uniform channel distribution, approximately 0.8% of the channels are expected to reside in this region; thus, values of 2–10% already represent substantial clustering, while values of 20–100% correspond to highly non-physiological distributions. Despite this, no EpC is observed in any of the considered cases. Even when the extracellular diffusion coefficients are reduced by a factor of *κ* = 6, EpC does not occur (upper right panel of Figure 12).

**Figure 12:**
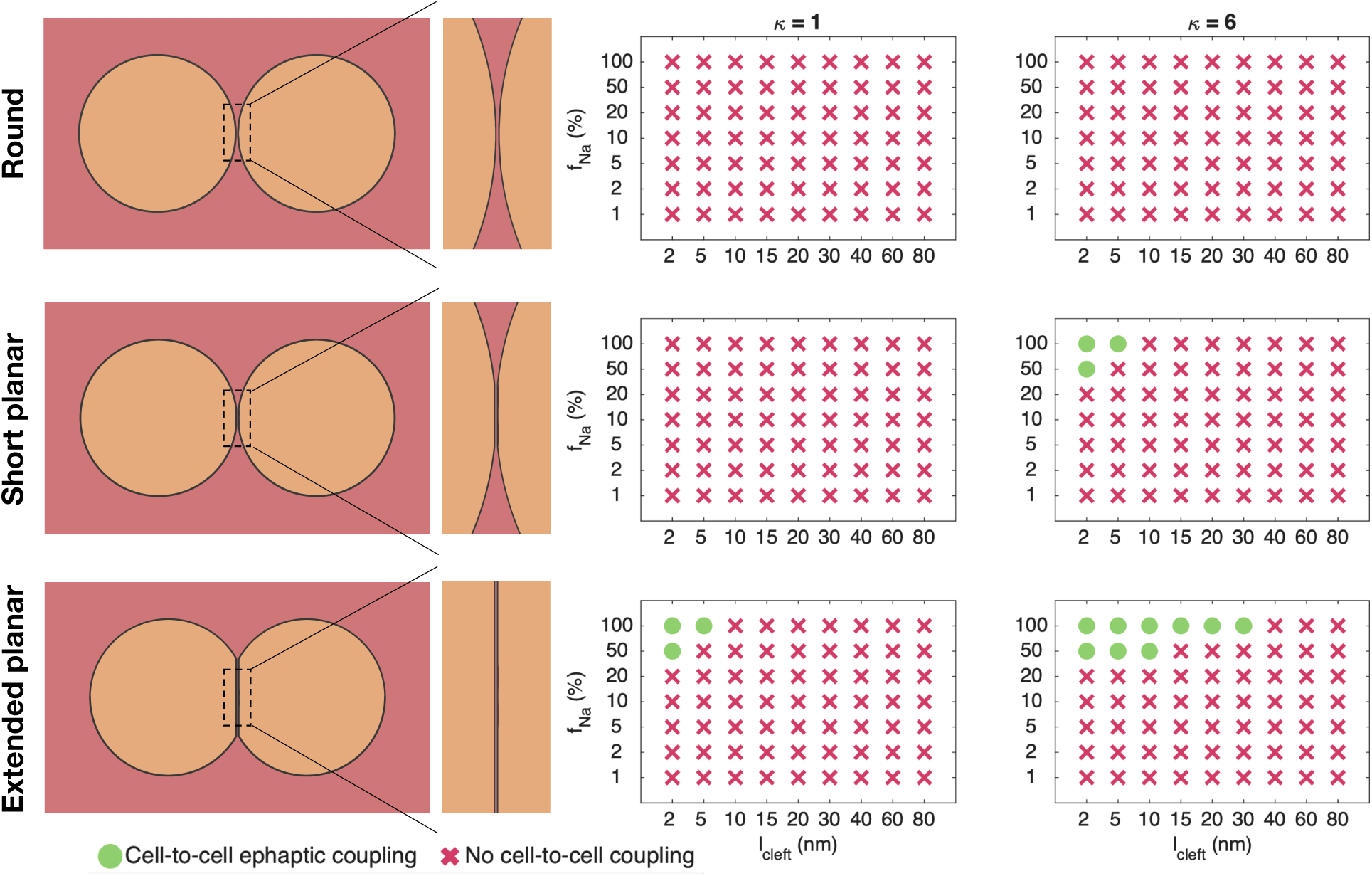
Occurrence of action potential transfer via ephaptic coupling in the side-by-side configuration. Here, *l*_cleft_ denotes the minimal distance between the cells, *f*_Na_ denotes the fraction of sodium channels localized to a 0.5 *µ*m-wide membrane patch at the region where the cells are closest, and *κ* denotes the factor by which the diffusion coefficients are divided in the extracellular space between the cells. We consider three different cell geometries: round cells, cells with a short (2 *µ*m) planar region facing the adjacent cell, and cells with an extended (10 *µ*m) planar region facing the adjacent cell. Details of the simulation setup and the criterion used to determine successful coupling are provided in the Methods section.

To assess the role of geometry, we also consider two progressively flattened cell shapes, shown in the middle and lower panels of Figure 12. For the short planar geometry, EpC is only observed for very small cell separations (2–5 nm), and only when diffusion is reduced and NaChs are highly clustered near the region of closest separation (above 50%). For the extended planar geometry, EpC becomes somewhat more likely, but still requires unrealistically high degrees of NaCh clustering in the same region.

## 3 Discussion

Our simulations identify geometrical and biophysical conditions under which ephaptic excitation transfer between cardiomyocytes may occur. In particular, they provide an explanation for the apparently opposing outcomes of side-by-side and end-to-end experiments, while also clarifying the roles of cleft width, NaCh organization, restricted diffusion, potassium channels, and GJ coupling.

### 3.1 Cell geometry reconciles apparently conflicting experiments

In [1], no electrical interaction or excitation transfer was observed when isolated cardiomyocytes were placed side by side, and this was interpreted as evidence against ephaptic transmission. In contrast, action potential transfer was later reported between isolated cardiomyocytes placed end to end in the absence of measurable GJ coupling, [2].

Our simulations are consistent with both observations. Ephaptic excitation transfer did not occur for round cells placed side by side, even with nanometer-scale separation and restricted extracellular diffusion. Transfer was only obtained after introducing flattened cell geometries and unrealistically strong localization of sodium channels near the region of closest contact. By contrast, excitation transfer occurred over a substantially wider range of conditions when the NaCh-rich membranes of the IDs faced each other in the end-to-end configuration.

The different experimental outcomes can therefore be explained by the geometrical configurations. End-to-end apposition provides a large, closely opposed membrane area with a high NaCh density, allowing sodium influx to generate a sufficiently negative extracellular potential to activate the neighboring cell. Side-by-side apposition involves a smaller region of close contact along the lateral membranes, where the NaCh density is expected to be lower.

This interpretation is consistent with the explanation proposed in [2], where it was argued that the earlier side-by-side experiments did not bring the NaCh-rich plicate regions of the ID into close apposition. Our simulations provide support for this explanation by showing that cell orientation and NaCh localization fundamentally alter the conditions for ephaptic excitation transfer.

### 3.2 Conditions for ephaptic excitation transfer

In our simulations, pure ephaptic excitation transfer required a sufficiently narrow extracellular cleft together with substantial localization of NaChs at the ID. As shown in Figure 3, transfer became more likely as the cleft width decreased and the fraction of NaChs located at the ID increased. Restricted ionic diffusion within the cleft further promoted transfer, as illustrated in Figure 7.

The spatial organization of the NaChs was also important. For a fixed fraction of channels at the ID, concentrating the channels into fewer, larger clusters strengthened EpC. The underlying mechanism is illustrated in Figures 4 and 5: prejunctional NaCh activation generates a negative extracellular potential in the cleft, which activates NaChs in the postjunctional membrane. This activation occurred more rapidly when the channels were concentrated into fewer clusters.

The dependence on cleft width was not monotone. For several parameter combinations, the activation delay was smallest at an intermediate cleft width, whereas both narrower and wider clefts produced longer delays, as shown in Figure 9. Beyond a critical width, excitation transfer failed abruptly rather than slowing gradually, indicating a threshold transition from successful coupling to block.

These results identify combinations of cleft width, NaCh localization, clustering, and diffusion restriction that permit pure EpC. They should be interpreted as qualitative indications: pure EpC is favored by increased NaCh localization at the ID, by concentration of NaChs into fewer clusters, and by an intermediate cleft width, rather than as precise quantitative estimates for intact cardiac tissue.

### 3.3 Effects of gap junctions and potassium channels

As shown in Figure 11, GJs had a much stronger effect on excitation transfer than pure EpC. Increasing GJ conductance substantially reduced the activation delay and increased the estimated conduction velocity. In addition, GJs replaced the abrupt transition from ephaptic conduction to block by a more gradual slowing of activation.

Localization of *I*_K1_ channels at the ID strongly altered the local potassium concentration but had only a modest effect on activation delay, as illustrated in Figure 10. This suggests that ID-localized potassium currents may influence the cleft environment without substantially changing the conditions for ephaptic excitation transfer.

### 3.4 Limitations

The end-to-end simulations include only two half-cells. This is sufficient for studying excitation transfer across a single ID, but several full cells would be required for reliable estimates of tissue-scale conduction velocity, particularly when GJs are present. The symmetric half-cell configuration may also favor EpC relative to a corresponding pair of full cells.

As in all numerical computations, spatial and temporal convergence must be considered. Mesh and time-step refinement are examined in Supplementary Figure S1, and the results indicate that the reported solutions are adequately converged. For computational efficiency, the end-to-end cells were represented as cuboids and solved using finite differences on highly adaptive meshes. We were unable to obtain a comparable degree of mesh adaptivity with the Gmsh-generated meshes used by the FEM/MFEM implementation. The side-by-side simulations were therefore restricted to two dimensions.

Finally, the cellular and ID geometries are strongly simplified. This allowed us to focus on the electrodiffusive dynamics described by the full PNP equations, but omits the tortuous structure, heterogeneous cleft widths, and detailed organization of ID nanodomains represented in more anatomically realistic models, [23].

## 4 Methods

In this section, the mathematical model used in our computations is described.

### 4.1 Model equations and parameters

We model the ionic concentrations ([*k*]) and the electrical potential (*φ*) in, between and around two neighboring cells using the Poisson-Nernst-Planck (PNP) model (see, e.g., [33, 34]) :

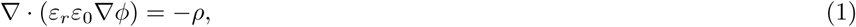

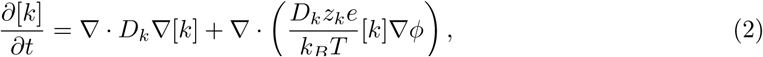

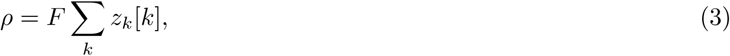

for *k* = *{*Na^+^, K^+^, Ca^2+^, Cl*^−^}*. Here, *ρ* is the charge density and *ε_r_*, *ε*_0_, *D_k_*, *z_k_*, *e*, *k_B_*, *T*, and *F* are parameters and physical constants and parameters. These are specified in Table 1. The initial conditions for the ionic concentrations are given in Table 2.

**Table 2:** Initial conditions for the ionic concentrations based on [38, 37, 39].

| Ion | Extracellular | Intracellular |
| --- | --- | --- |
| $\text{Na}^+$ | 100 mM | 12 mM |
| $\text{K}^+$ | 5.4 mM | 125 mM |
| $\text{Ca}^{2+}$ | 1.4 mM | 0.0001 mM |
| $\text{Cl}^-$ | 108.2 mM | 15 mM |

### 4.2 Model geometry

The model geometry differs between the end-to-end and side-by-side configurations, as described below.

#### 4.2.1 End-to-end configuration

In the end-to-end configuration, we consider two cardiomyocytes represented as rectangular cuboids (see Figure 1). The width of the cells is 18 *µ*m, and we include half of the cells in the longitudinal direction (50 *µ*m each). The cell membrane separating the intracellular and extracellular domains is modeled with a thickness of 5 nm. The cells are separated by an extracellular cleft with width varying between 5 and 80 nm, and are surrounded by a bulk extracellular bath of width 1 *µ*m.

We impose no-flux Neumann boundary conditions in the longitudinal (*x*) direction and Dirichlet boundary conditions on the remaining boundaries in the *y*- and *z*-directions for both the electric potential, *φ*, and ionic concentrations. The Dirichlet boundary conditions are given by *φ* = 0 mV and the extracellular ionic concentrations as specified in Table 2.

#### 4.2.2 Side-by-side configuration

In the side-by-side configuration, the cells are modeled as cylinders, and we assume translational symmetry in the longitudinal direction, reducing the computational domain to two dimensions (see Figure 2). Each cell has a diameter of 20 *µ*m, and the membrane thickness is 5 nm. Dirichlet boundary conditions are applied on the outer boundary of the extracellular domain, as in the end-to-end configuration. The minimal distance between the cells and the outer boundary of the extracellular space is 1 *µ*m.

### 4.3 Transmembrane ionic fluxes

In this section, the transmembrane ionic fluxes is described.

#### 4.3.1 Ion channel fluxes

The ion channel fluxes used in our simulations are based on the human ventricular action potential model of Grandi et al. [46]. We include the inward rectifier potassium current, *I*_K1_, which is required to maintain a physiological resting membrane potential, as well as the three currents whose peak current densities exceed 1% of the peak fast sodium current density during the action potential upstroke in the Grandi et al. model: the fast sodium current, *I*_Na_, the L-type calcium current, *I*_CaL_, and the transient outward potassium current, *I*_to_.

Ion channel fluxes are implemented as internal boundary conditions on the intracellular and extracellular sides of the membrane, following the approach described in [25]. The flux through channels of type *j* carrying ion species *k* within a given cluster or membrane patch is given by

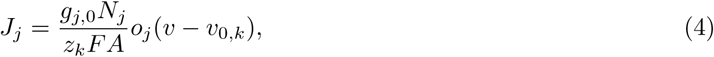

where *g_j,_*_0_ is the single-channel conductance, *N_j_* is the number of channels in a cluster or membrane patch of area *A*, *z_k_* is the valence of ion species *k*, and *F* is Faraday’s constant. Furthermore, *v* = *φ_i_ − φ_e_* is the membrane potential and *v*_0*,k*_ is the Nernst equilibrium potential,

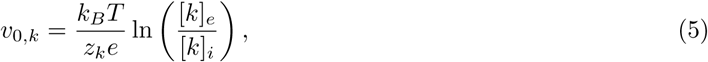

where [*k*]*_e_* and [*k*]*_i_* denote the extracellular and intracellular concentrations of ion species *k*, respectively. The channel open probability, *o_j_*, is described by the gating models given in the Grandi et al. model [46]. The quantities *o_j_*, *v*, and *v*_0*,k*_ are evaluated and applied locally on each mesh element belonging to the relevant cluster or membrane patch.

The total number of channels of each type is estimated from the whole-cell conductance densites, *G_j_*, reported by Grandi et al. [46]. Specifically, we multiply the conductance densities by the membrane area assumed in that model (*≈* 13, 000 *µ*m^2^) and divide by the corresponding single-channel conductances reported in the literature. The resulting channel numbers are listed in Table 3. For the end-to-end configuration, which represents half-cells, the number of channels is reduced by a factor of two. In the two-dimensional side-by-side configuration, the factor 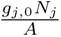 is replaced directly by the conductance density *G_j_* from the Grandi et al. model.

**Table 3:** Ion channels, single channel conductances and total number of channels of each type. The number of channels are estimated from [46], and the single channel conductances are based on the provided references.

| Channel, $j$ | Ionic species, $k$ | $g_{j,0}$ | $N_j^{\text{tot}}$ | Ref. |
| --- | --- | --- | --- | --- |
| Na | Na<sup>+</sup> | 20 pS | 144500 | [40, 41, 42] |
| K1 | K<sup>+</sup> | 5 pS | 8800 | [43] |
| to | K<sup>+</sup> | 7.5 pS | 1920 | [44] |
| CaL | Ca<sup>2+</sup> | 8 pS | 2000 | [45] |

#### 4.3.2 Channel clustering

For the side-by-side configuration and the lateral membranes in the end-to-end configuration, the ion channels are assumed to be uniformly distributed throughout the membrane patches. All ion channels are assumed to be located here, except for a certain percentage of the *I*_Na_ or *I*_K1_ channels, which are assumed to be located at the ID. At the ID, these channels are organized into *N_c_*clusters. The number of channels in each cluster is given by

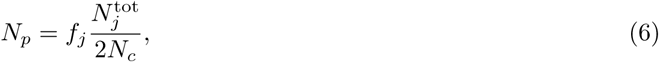

where *f_j_* denotes the fraction of channels of type *j* located at the ID, and *N* ^tot^ is the total number of channels of type *j* in a full cell. The factor 2 in the denominator accounts for the fact that only half of each cell is included in the end-to-end simulations. Within a cluster, each channel is assumed to occupy an area of 10 nm *×* 10 nm. The clusters are assumed to be square, with side length *w_c_* determined by the number of channels in the cluster:

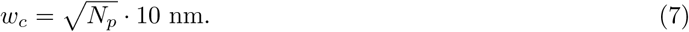

Accordingly, each cluster occupies an area of *w_c_ × w_c_*.

#### 4.3.3 Gap junction flux

To investigate the effect of EpC in the presence of GJs, we distribute a GJ conductance, *G*_gap_, uniformly over the ID. The GJ current is given by

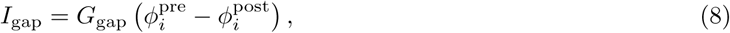

where *G*_gap_ is the gap junction conductance (in zS) and *φ_i_*^pre^ and *φ_i_*^post^ denote the intracellular potentials (mV) adjacent to the ID membrane in the prejunctional and postjunctional cells, respectively. The total gap junction current is partitioned among the ionic species according to their relative transport numbers [40],

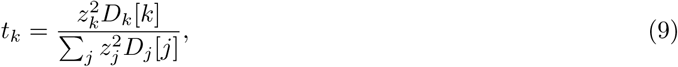

where [*k*] denotes the average concentration of ion species *k* on the prejunctional and postjunctional sides of the gap junction. The corresponding flux of ion species *k* is then given by

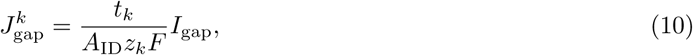

which is imposed as an internal boundary condition on the intracellular sides of the prejunctional and postjunctional ID membranes. Here, *A*_ID_ is the area of the ID (in nm^2^).

### 4.4 Simulation setup

In our investigations of EpC, we run simulations in three steps.

#### Part I: Initialization of the resting state

The first step computes the resting state of the cells. Simulations are initialized from the electroneutral conditions given in Table 2, with all *I*_K1_ channels fully open (*o*_K1_ = 1) and all other ion channels closed. The simulation is run until an equilibrium solution is reached, representing the resting state of the cells, following the procedure in [18, 25].

#### Part II: Initial transfer simulation

The second simulation step starts from the resting state solution obtained in Part I. In this step, the ion channel open probabilities are governed by the gating models described in Section 4.3.1. An action potential is initiated in the prejunctional cell by prescribing a time-dependent Dirichlet boundary condition. In the end-to-end configuration, this boundary condition is applied over the central 10 *µ*m *×* 10 *µ*m of the left boundary, whereas in the side-by-side configuration it is applied over a circular region of diameter 1 *µ*m near the left side of the prejunctional cell. The prescribed waveform is taken from the initial phase of an action potential generated by the Grandi et al. membrane model [46].

#### Part III: Secondary transfer simulation

The third step is identical to Part II, except that the Dirichlet boundary condition is taken from the intracellular potential recorded near the right side of the postjunctional cell during the preceding simulation. This is done to represent propagation of the action potential from the postjunctional cell to a third downstream cell.

### 4.5 Definition of cell-to-cell coupling, activation delay, and conduction velocity

For the end-to-end configuration, we define a prejunctional and a postjunctional observation point at the centers of the leftmost and rightmost boundaries, respectively, representing the centers of the prejunctional and postjunctional cardiomyocytes. Cell-to-cell coupling is considered to occur if the intracellular potential at the postjunctional observation point exceeds *−*30 mV during the simulation. The activation delay, *τ*, is defined as the time difference between the moments at which the intracellular potential at the prejunctional and postjunctional observation points first exceeds *−*30 mV. The conduction velocity (CV) is then defined as the distance between the cell centers (100 *µ*m) divided by *τ*.

For the side-by-side configuration, cell-to-cell coupling is considered to occur if the intracellular potential in an observation point near the right end of the postjunctional cell exceeds *−*30 mV during the simulation.

### 4.6 Numerical methods

We solve the PNP system numerically using the coupled scheme from [25, 27], which enables large time steps. For the Part I simulations, a time step of 0.3 ms is used, while for the Part II and Part III simulations a smaller time step of 0.025 ms is employed to capture the AP upstroke dynamics. Refinement of the temporal and spatial discretization parameters is investigated in Supplementary Figure S1.

The PNP equations and the ion channel gating dynamics are treated separately using a standard operator splitting approach (see, e.g., [34]). The gating variables are updated using an explicit forward Euler scheme with a local time step of 0.0001 ms.

For the 3D end-to-end configuration, the PNP system is discretized using a finite difference method on an adaptive mesh (see, e.g., [18]). For the 2D side-by-side configuration, the PNP equations are solved using a standard Galerkin finite element method implemented in the MFEM C++ library [47, 48], employing first-order *H*^1^ elements for both the electric potential and ionic concentrations. The computational finite element meshes are generated using Gmsh [49].

## Supporting Information

The Supporting Information provides a numerical convergence study examining the effects of temporal and spatial resolution on the simulated extracellular potential.

**Figure S1:**
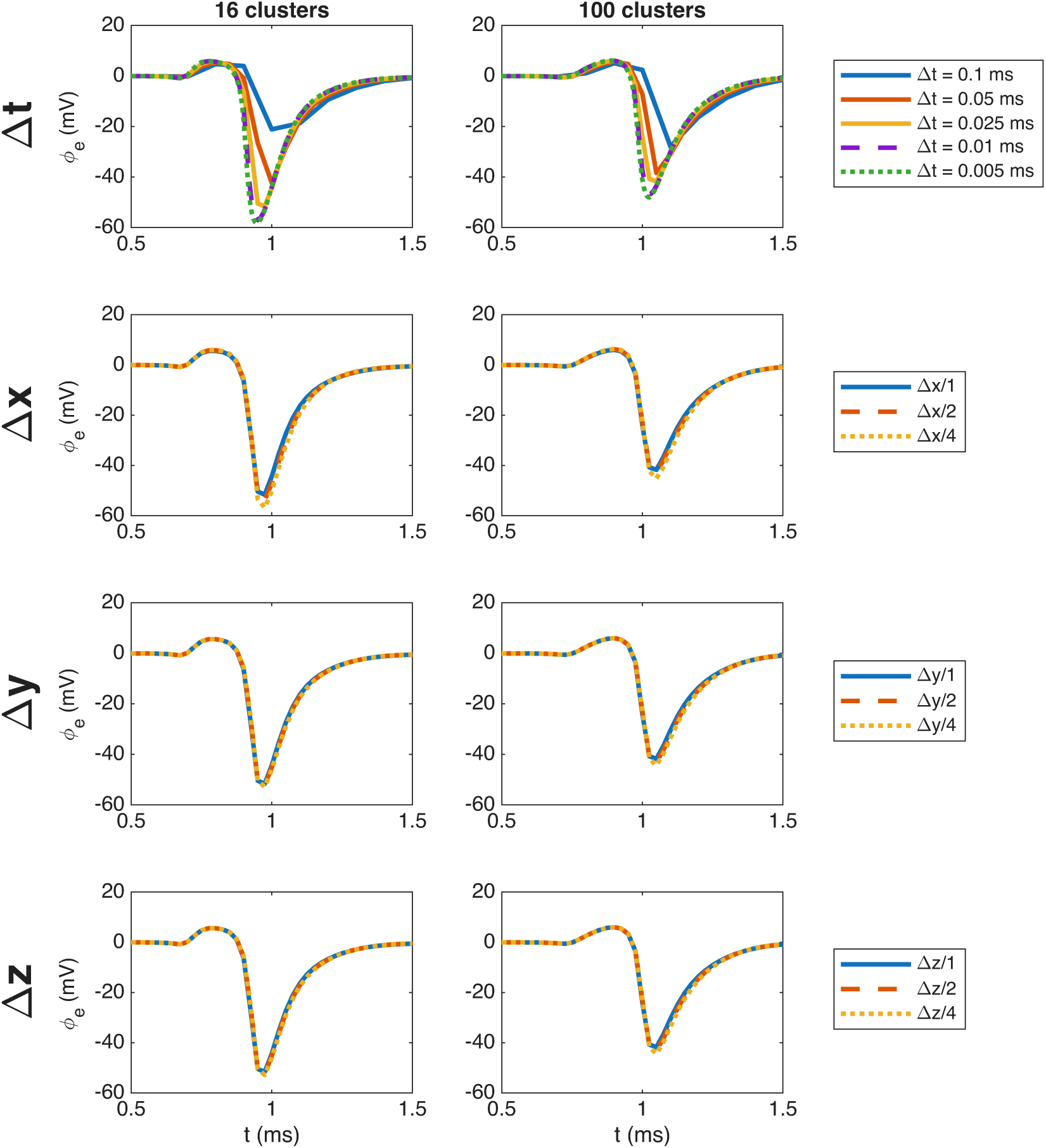
Effect of the temporal and spatial resolution on the simulated extracellular potential. To reduce the computational cost of the refinement study, only the prejunctional cell is included. The extracellular potential is evaluated at the center of the extracellular boundary opposite the cell end, 15 nm from the membrane. A no-flux (Neumann) boundary condition is imposed on this boundary. Temporal resolution is investigated by performing simulations with five different time steps using the default adaptive mesh. Spatial resolution is investigated by considering two successive levels of refinement of the adaptive mesh (see, e.g., [18]) in each spatial direction while keeping the default time step fixed. In the default adaptive mesh, the Δ*x* values are in the range 0.5 nm to 12.5 *µ*m, and Δ*y* and Δ*z* are in the range 5 nm to 1.5 *µ*m.

